# Loss of CARM1 alters the developmental programming of Glioma stem-like cells and creates a druggable NGFR/NTRK dependency

**DOI:** 10.1101/2025.04.11.647869

**Authors:** Dejauwne L. Young, Jennifer Aguilan, Ronald Cutler, Stephanie Stransky, Joseph D. DeAngelo, Jacob S. Roth, Beata Malachowska, Justin Vercellino, Brett I. Bell, David Shechter, Philip J. Tofilon, Richard E. Phillips, Chandan Guha, Simone Sidoli

## Abstract

A key driver of Glioblastoma (GBM) heterogeneity and therapy resistance is the capacity of glioma stem-like cells (GSCs) to hijack developmental signaling programs. However, it remains unclear how GSCs regulate these adapted developmental signaling pathways and how these pathways might be therapeutically exploited. The arginine methyltransferase, CARM1, has been shown to play critical roles in maintaining stem cell pluripotency, preventing differentiation, and recently was discovered to be upregulated in Glioblastoma. To date, there is little to no understanding of the role that CARM1 plays in regulating developmental processes in Glioblastoma. To address this gap in knowledge, we applied a multi-omics approach to characterize developmental processes that are specifically regulated by CARM1 in GSCs. We found that loss of CARM1 results in dysregulation of several developmental markers: ARX, GFAP, NGFR, PDGFRA and results in both a proteomic and transcriptomic shift towards the radial glia cell lineage. Moreover, CARM1 depleted cells reprogram their signaling to develop an increased survival dependency on NGFR/NTRK signaling and are hypersensitive to the FDA approved brain penetrant NTRK inhibitor—Entrectinib. Mechanistically, we find that NFIA is a CARM1 substrate and can repress NGFR signaling just as CARM1 does, and thus the CARM1/NFIA relationship is likely a key regulator of NGFR/NTRK signaling in GSCs. Altogether, we demonstrate that CARM1 regulates the cell lineage of GSCs at the transcriptomic and proteomic level, and naturally represses NGFR/NTRK signaling—likely through CARM1 dependent methylation of NFIA. Further, CARM1 depletion leads GSCs to develop a survival dependency on NGFR/NTRK signaling and creates a therapeutic vulnerability to NTRK inhibition.

## Introduction

Glioblastoma (GBM) is the most common malignant primary brain tumor and is universally fatal, with an average survival of 14-16 months from initial diagnosis [1, 2]. GBM aggressiveness is due to a complex interplay of tumor heterogeneity, chemo-radiation resistance, and tumor invasiveness largely driven by Glioblastoma stem-like cells (GSCs) [3]. GSCs are characterized by their expression of multiple stem cell markers, capacity to self-renew and differentiate, along with their high tumorigenicity in-vivo [4]. Moreover, GSCs rely heavily on epigenetic regulation to maintain their stem-like character and evade therapy [5–7].

One key regulator of epigenetics and transcription in mammals is protein arginine methylation. Arginine methylation is a vital post-translational modification occurring in mammals and are catalyzed by either type I protein arginine methyltransferases (PRMTs) or type II PRMTs [8].

Type I PRMTs include PRMT1/2/3/4/6/8 which are responsible for catalyzing asymmetric dimethylarginine (ADMA) and type II PRMTs including PRMT5/9 which catalyze symmetric dimethylarginine (SDMA) [9]. Arginine methylation often functions to promote protein-protein or protein-nucleic acid interactions, and when lost can lead to alterations in gene expression or protein/cellular functions [10].

Arginine methylation specifically plays a critical role in glioblastoma therapy resistance and chromatin regulation. It has been shown that PRMT1 and PRMT5 are upregulated in GBM and can regulate vital processes such as proliferation, splicing, senescence, and stemness [8, 11–13]. A member of the Type I PRMT family that remains under-investigated in GBM is Coactivator-Associated Arginine Methyltransferase (CARM1)/PRMT4. CARM1 specifically deposits ADMA marks on histones and chromatin modulating proteins to exert regulation on transcription and RNA processing [14]. During embryogenesis, CARM1 has been shown to be involved in regulating cell fate decisions [15, 16]. However, how CARM1 alters developmental signaling and cell fate in Glioblastoma remains unknown. Since CARM1 is overexpressed in GBM [17] and is implicated in normal tissue development, targeting CARM1 may be a promising strategy to selectively target the GSC population. To date, it remains unknown how CARM1 impacts GSC character and to what degree these changes might be therapeutically exploited.

Using CRISPR/Cas9 and a multi-omics approach, we show that loss of CARM1 alters the growth of GSCs and results in widespread changes in various histone post-translational modifications (PTMs) often involved in development, beyond H3R17. We also find that CARM1 regulates developmental signaling changes on both a transcriptomic and proteomic level and represses the radial glial cell lineage in GSCs. Moreover, we observe that loss of CARM1 drives up NGFR/NTRK signaling in GSCs, likely through methylation of its substrate NFIA, and this new dependency can be targeted with NTRK inhibitors like Entrectinib. Overall, our work provides novel insight on how CARM1 regulates cell fate commitment in GSCs and reveals a new therapeutic approach for the treatment of Glioblastoma.

## Material and methods

### Cell culture

Glioma stem-like cell lines (GSCs) NSC11 and NSC20 are two patient derived cell lines provided as frozen stocks by Dr. Frederick Lang, MD Anderson Cancer Center, Houston, TX. GSCs were grown as neurospheres in stem cell media (DMEM/F12, 50ng/ml EGF and bFGF, 2% B27 without vitamin A, and 1% penicillin-streptomycin) and CD133+ cells were selected by FACS as previously reported [18]. All cell lines were maintained in culture (5% O_2_ and 5% CO_2_ incubators) for less than two months from thawing and confirmed to be negative for mycoplasma with PCR. GSCs cell morphology and growth was constantly monitored for health, and NSC11 and NSC20 cell line differences were authenticated using proteomics.

### Lentiviral CRISPR/Cas9

We used a two plasmid approach to ablate target genes. The lentiCas9-Blast (plasmid #52962) and pXPR_050 (plasmid #96925) was purchased from addgene. Custom sgRNA sequences were made with CRISPick and cloned as described previously [19]. Lentiviral particles were produced in HEK293T cells by calcium phosphate transfection. Lentivirus was next concentrated using polyethylene glycol (systems biosciences LV810A-1) and stored in cryotubes in -80°C. Transduction of GSCs was performed by mixing 25ul of concentrated lentivirus with 1 x 10^6^ GSCs in 100ul of stem cell media and incubated for 2 hours at 5% O_2_ and 5% CO_2_. After lentiviral infection GSCs were cultured for 5 days in normal stem cell media, then selected for 8 days with 10ug/ml of blasticidin (ThermoFisher A1113903) and/or 4ug/ml of puromycin (Sigma P9620). Gene depletion was then validated with western blot.

sgCARM1-1: sense: 5’-AGCACGGAAAATCTACGCGG-3’ antisense: 3’-CCGCGTAGATTTTCCGTGCT-5’

sgCARM1-2: sense: 5’-TGGAGCACGGAAAATCTACG-3’ antisense: 3’-CGTAGATTTTCCGTGCTCCA-5’

### siRNA knockdown

We used lipofectamine RNAiMAX transfection reagent (ThermoFisher) to generate siRNA knockdowns. We purchased two different siRNA against NFIA (ThermoFisher s9477, s9478) and one against NGFR (ThermoFisher s9540). We followed the RNAiMAX protocol for transfection. In short, 1 x 10^6^ GSCs were seeded into 10ml of stem cell media with 40nM of siRNA-lipofectamine complexes against siRNA of target gene or siCTRL (negative control) and placed in incubator for 24 hours. GSCs were then gently collected, spun down, and reseeded with fresh media without siRNA and placed in incubator for another 4 days. Fresh media was added accordingly. siRNA sequences are below:

siNFIA-1: sense: 5’-CGAAAACGAAAAUACUUCAtt-3’ antisense: 5’-UGAAGUAUUUUCGUUUUCGgg-3’

siNFIA-2: sense: 5’-CACAUCACCCAUUAUCCAtt-3’ antisense: 5’-UGGAUAAUGGGUGAUGUCGgg-3’

siNGFR: sense: 5’-CCACAGCAGGUGUCAUAUAtt-3’ antisense: 5’-UAUAUGACACCUCUGUGGtg-3’

### Western blot

Fifteen to twenty micrograms (lysed with RIPA buffer and measured with BCA, ThermoFisher) of cell lysates were mixed with 2-4x Laemmli Sample Buffer and β mercaptoethanol, then ran down SDS-PAGE 4-15% gradient gel and transferred to a PVDF membrane. The membrane was blocked with 5% non-fat dry milk in Tris-Buffered Saline (TBS)-Tween 0.1% for 1 hour. The membrane was then incubated with blocking buffer and various antibodies: CARM1 (ThermoFisher A300-421A 1:2000), NGFR (Cell Signaling #8238 1:1000), ERK1/2 (Cell Signaling #9102 1:1000), phospho-ERK1/2 (Cell Signaling #4370 1:1000), NFIA (Cell Signaling #69375 1:1000), AKT (Cell Signaling #4691 1:1000), phospho-AKT (Cell Signaling #9271 1:1000), GAPDH (Santa Cruz #47724 1:1000).

### Cell viability and drug treatments

For growth curves, 1.5 x 10^3^ GSCs were seeded in 100ul of stem cell media in opaque 96 well plates, growth was measured on day 0, 3, and 6 with Cell Titer-Glo 2.0 (Promega G9242). For drug treatments, 5 x 10^3^ GSCs were seeded into 100ul of stem cell media with various dosages of either Entrectinib (MedChemExpress HY-12678) or Capivasertib (MedchemExpress HY-15431) for 96 hours and compared to DMSO vehicle controls. Cell viability was then measured with Cell Titer-Glo 2.0. Drug and growth curves were plotted with Prism (version 10) for visualization, data is reported as mean +/- SEM. Student’s T-test was applied for statistical significance.

### Clonogenic Assays

For clonogenic assays [20], GSCs were dissociated into single cells and 100-500 cells were seeded onto poly-l-lysine (sigma) coated 6 well plates in stem cell media and allowed to adhere overnight. Media was then removed and replaced with media containing 3uM of Entrectinib or DMSO and incubated for 72 hours. Media was then replaced with drug free media and colonies were allowed to form for 16 days. Next, colonies were stained and fixed with 0.5% crystal violet in 10% neutral buffered formalin. The number of colonies containing ≥ 50 cells were counted for sgSCR, sgCARM1-1, and sgCARM1-2. Entrectinib colony formation efficiency was normalized to DMSO colony formation efficiency for each sgRNA to account for differences in plating efficiencies. Data is reported as mean +/-standard deviation. The Student’s t-test was performed for statistical evaluation with alpha level of 0.05 for statistically significant results.

### Proteomics sample preparation

Cell pellets were prepped and digested based on S-trap proteomic filters (Protifi) based on the manufacturer’s instructions. Briefly, dry cell pellets were resuspended with 50ul of 5% SDS and 1ul of Pierce universal nuclease (ThermoFisher 88701), followed by 1 hour incubation in DTT (dithiothreitol, Sigma) at 37°C, and a 30 minute incubation in 20mM iodoacetamide (Sigma). Next, phosphoric acid was to samples at a concentration of 1.2% and proteins were diluted in S-trap binding buffer (90% methanol and 10mM ammonium bicarbonate), then added to S-trap filters. Samples were spun down for 1 minute at 500G, then filters were washed three times with S-trap binding buffer. One 1ug of sequence grade trypsin (Promega) or chymotrypsin (Promega) was mixed with 50mM ammonium bicarbonate and added to samples in S-trap filters. Proteins were then digested overnight at 37°C, then sequentially eluted with 60ul of 50mM ammonium bicarbonate, 0.1% trifluoroacetic acid (TFA), and 60% acetonitrile/0.1% TFA. Peptides were then completely dried in a speed vacuum and stored in -80°C freezer until desalting or other purposes such as CARM1 substrate identification or phospho-proteomics.

For identification of CARM1 asymmetric di-methyl arginine (ADMA) substrates we used the PTMScan kit (Cell Signaling #18303S) as previously [21], and largely followed the kit instructions. Briefly, after drying of S-trap eluted peptides as described above, peptides were resuspended in PTMScan IAP buffer (10% of samples were saved for analysis of total proteome, 90% used for PTMScan) and mixed with appropriate volume of ADMA antibodies bound to magnetic beads and rotated overnight at 4°C. After several washes peptides were eluted off magnetic beads with IAP Elution Buffer (0.15% TFA + 99.85% water), dried in speed vacuum, and stored in -80°C freezer.

For identification of phospho-peptides we used a previous protocol as described here [22]. In short, after elution and drying of digested peptides, peptides were resuspended in 100ul of 0.1% TFA and 10% of sample was taken for total proteome analysis, remaining 90% proceeded with phosphorylation pulldown. We added 0.6 mg of titanium beads (GL Sciences #5010-21315) per 100 μ of protein and incubated for 10 minutes with shaking at 1400 rpm, removed supernatant carefully. We repeated this process one more time to capture all phospho-peptides. After several washes phospho-peptides were eluted with Elution buffer (80ul of Ammonium Hydroxide + 980ul HPLC water, pH = 11.3), dried in speed vacuum, and stored in -80°C freezer.

### Histone sample preparation

Histones were extracted as previously described [23, 24]. Dry cell pellets were dissolved in 350ul of 0.2 M sulfuric acid and incubated for 4 hours at 4°C with rotation. Samples were mixed with 150ul (or 33%) trichloroacetic acid (TCA) overnight at 4°C. Then samples were spun down for 3400G for 5 minutes, and the supernatant was then disposed of. Next histones were sequentially washed with 0.1 HCL in cold acetone and 100% cold acetone, then air dried. Histones were then resuspended in 50ul of 50mM ammonium bicarbonate with 20% acetonitrile. In the fumigation hood, histones were derivatized with addition of 2ul of propionic anhydride and 10ul of ammonium hydroxide were added and incubated for 10 minutes, these steps were then repeated once again and samples were then dried in a speed vacuum. Next, 1ug of sequencing grade trypsin was diluted in 50mM Ammonium Bicarbonate and incubated overnight at 37°C then derivatized again the following day. Samples were then dried in a speed vacuum and stored in -80C freezer.

### Sample desalting

Before mass spectrometry analysis samples were desalted with 1mg of Oasis HLB C-18 resin (Waters) in 96-well filter plates (Orochem). Briefly, plates were loaded with HLB C-18 resin, then washed, and samples resuspended in 0.1% TFA were added to plates. Samples were washed with 0.1% TFA, then eluted with 0.1 TFA/60% Acetonitrile, and dried in a speed vacuum.

### Gene Set Enrichment Analysis

Gene Set Enrichment Analysis was performed using the R (v4.3.1) package clusterProfiler [25]. Gene sets used were from MSigDB C2 collection and Gene Ontology. For proteomics data, we averaged the log2foldchange and -log_10_p-value of sgCARM1-1 and sgCARM1-2 then the data was rank-ordered by (sign (log2foldchange) * -log_10_p-value). For RNA-seq the data was also rank-ordered by (sign (log2foldchange) * -log_10_adjp-value).

### TCGA analysis

To access TCGA data of CARM1 expression in lower grade glioma and GBM patients and the survival of GBM patients split by this expression, we used UCSC Xena browser [26]. Data analysis was performed with R (v.4.3.1) with survival and survminer libraries.

### Proteomic analysis

After desalting, samples were resuspended in 0.1% Formic Acid (FA) and loaded onto the Bruker nanoElute 2 coupled online with the timsTOF HT (Bruker). Chromatography separation was used using a two-column system consisting of a C18 trap cartridge (ThermoFisher #174500) and an analytical PepSep XTREME column of 25cm x 150µm x 1.5µm (Bruker #1893476). Samples were separated using a 45 minute gradient from 4 to 30% buffer B at a flow rate of 700nl (Buffer A: 0.1% FA, Buffer B: 80% acetonitrile + 0.1% FA). The spectra were acquired in dia-PASEF in the range of 100-1700 m/z and 0.6 1/K0 [V-s/cm-2] to 1.6 1/K0 [V-s/cm-2]. Ramp and accumulation times were set to 100. 32 DIA isolation windows were used in the range of 400-1201 m/z and 0.6 1/K0 [V-s/cm-2] to 1.6 1/K0 [V-s/cm-2]. The collision energy was ramped as a function of increasing mobility starting from 20 eV at 0.60 1/K0 to 59 eV at 1.6 1/K0. The MS raw data was searched using FragPipe version 22 [27] with the preloaded workflow “DIA_ SpecLib_Quant_diaPASEF”. Our digestive enzymes were trypsin or chymotrypsin (for CARM1 substrate analysis only). Using R, raw protein abundances were log2 transformed, normalized by median centering, and imputed using a normal distribution two standard deviations beneath the mean as described previously [28]. For CARM1 ADMA substrate and phospho-proteomics analysis we added another normalization step in which normalized ADMA peptides and phospho-peptides were divided by the normalized total protein abundance of their respective proteins. This approach allows us to quantify the differential abundance of the specific PTMs while removing any bias that may be introduced due to differences in total protein abundance. For statistical analysis, the F-test was performed to assess for protein/peptide variation followed by the appropriate heteroscedastic or homoscedastic t-test with alpha level of 0.05 for statistically significant results. Proteins/peptides were considered differentially expressed only if they were statistically significant in both sgCARM1-1 and sgCARM1-2.

### Histone post-translational modification (PTMs) analysis

Post desalting, samples were resuspended in 0.1% TFA and placed onto the Dionex RSLC Ultimate 300 and analyzed by an Orbitrap Fusion Lumos (ThermoFisher). A two column system was used to separate the histone peptides composed of a C18 trap cartridge (300 µm ID, 5 mm length) and an analytical column (75 µm, 25 cm). The analytical column was packed in our lab with reversed-phase Repro-Sil Pur C18-AQ 3 µm resin. Histone peptides were separated using a 30 minute gradient and at a flow rate of 300nl/min in DIA mode. The full MS scan was set to 300-1100 m/z in the orbitrap with a resolution of 120,000 (at 200m/z).

Raw data files were searched through EpiProfile 2.0 software [29], and the area under the curve was generated for each identified histone peptide. To properly normalize the data, the sum of every identified combination for a histone peptide was summed (total peptide abundance) and considered to be 100% and the area of each individual peptide was divided by the total peptide abundance to generate a ratio. The relative ratio of two isobaric forms was calculated by averaging the ratio for each fragment ion between the two. Using R, the final peptide table was log2 transformed, log2foldchange was calculated, and statistical testing was done first with the F-test followed by the appropriate heteroscedastic or homoscedastic t-test with alpha level of 0.05 for statistically significant results. All histone PTMs were only considered differentially expressed if they were statistically significant in both sgCARM1-1 and sgCARM1-2.

### Bulk mRNA transcriptome differential expression analysis

RNA was extracted from cell pellets using the RNeasy kit (Qiagen), quantified using a Qubit kit (Thermo Fisher Scientific) and high purity was confirmed by 260/280 and 260/230 ratios using a Nanodrop (Thermo Fisher Scientific). Libraries were prepared for mRNA analysis and sequenced on an Illumina Novaseq X using 150bp paired-end mode by Novogene Corp Inc., CA. Raw sequence reads were processed using the nf-core rnaseq pipeline (https://nf-co.re/rnaseq/3.14.0) using the star_rsem alignment option and min_mapped_reads set to 5 to align the reads to the GRCh38 primary genome assembly and gene level counts were obtained using the gencode v42 primary assembly annotation [30]. Gene filtering was performed using the zPKM program [31]. Correlation analysis of TPM values between replicate samples was performed to ensure high concordance. Since there were no significant differences between sgCARM1-1 and sgCARM1-2 we combined these replicates and compared them to sgSCR. Count normalization and differential expression testing was performed using DESeq2 and genes were considered significantly enriched if the adjusted p-value < 0.05.

### Qiagen Ingenuity Pathway Analysis (IPA) and over representation analysis

For analysis of only differentially expressed genes we used two tools, Gene Ontology over representation analysis (ORA) and Qiagen IPA. To identify the Gene Ontology molecular functions of CARM1 substrates we used clusterProfiler in R. We used Benjamini-Hochberg for multiple testing correction, only considered GO terms with an adjusted p-value < 0.05, and used all the identified proteins in the dataset as our background. For phospho-proteomic analysis we used Qiagen’s Ingenuity Pathway Analysis (IPA) software [32]. We uploaded our table of differentially expressed phospho-peptides including the phospho-sites to IPA and selected the default phosphorylation analysis as our core analysis. We downloaded tables of most enriched canonical pathways and upstream regulators. For upstream regulators we filtered out chemical drug/reagents/toxicants as this was not relevant to our work and sorted by descending bias-corrected z-score, only including regulators with p-value < 0.05. For canonical pathways, we filtered for pathways with a p-value < 0.05 and an absolute Z-score of 2 or greater, then sorted in descending order based on Z-score. Qiagen IPA results were visually represented with dot plots from R using the ggplot2 library.

## Results

### CARM1 predicts Glioma patient survival and regulates Glioma stem-like cell proliferation

We first wanted to understand the clinical impact of CARM1 in Glioblastoma, so we initially analyzed publicly available datasets from the TCGA. We identified that CARM1 is significantly overexpressed in high grade glioma in comparison to low grade glioma (**Fig. 1A**), in line with previous findings [17]. CARM1 mRNA expression is also strongly correlated with glioblastoma patient survival, with lower CARM1 expression correlating with increased patient survival (**Fig. 1B**). Since low CARM1 expression is associated with increased GBM patient survival and has been previously reported to slow cell growth in LN229 cells [33], we hypothesized that depletion of CARM1 would also slow proliferation of Glioma stem-like cells. As a model system, we used CRISPR/Cas9 along with a negative selection approach to ablate the CARM1 gene in CD133+ patient derived glioma stem-like cells (GSCs). We validate CARM1 depletion in two distinct GSCs using western blot (**Fig. 1C-D**). We tested GSC cell growth across multiple time points and found that loss of CARM1 slowed GSC cell growth in both cell lines (**Fig. 1E-F**).

**Figure 1:**
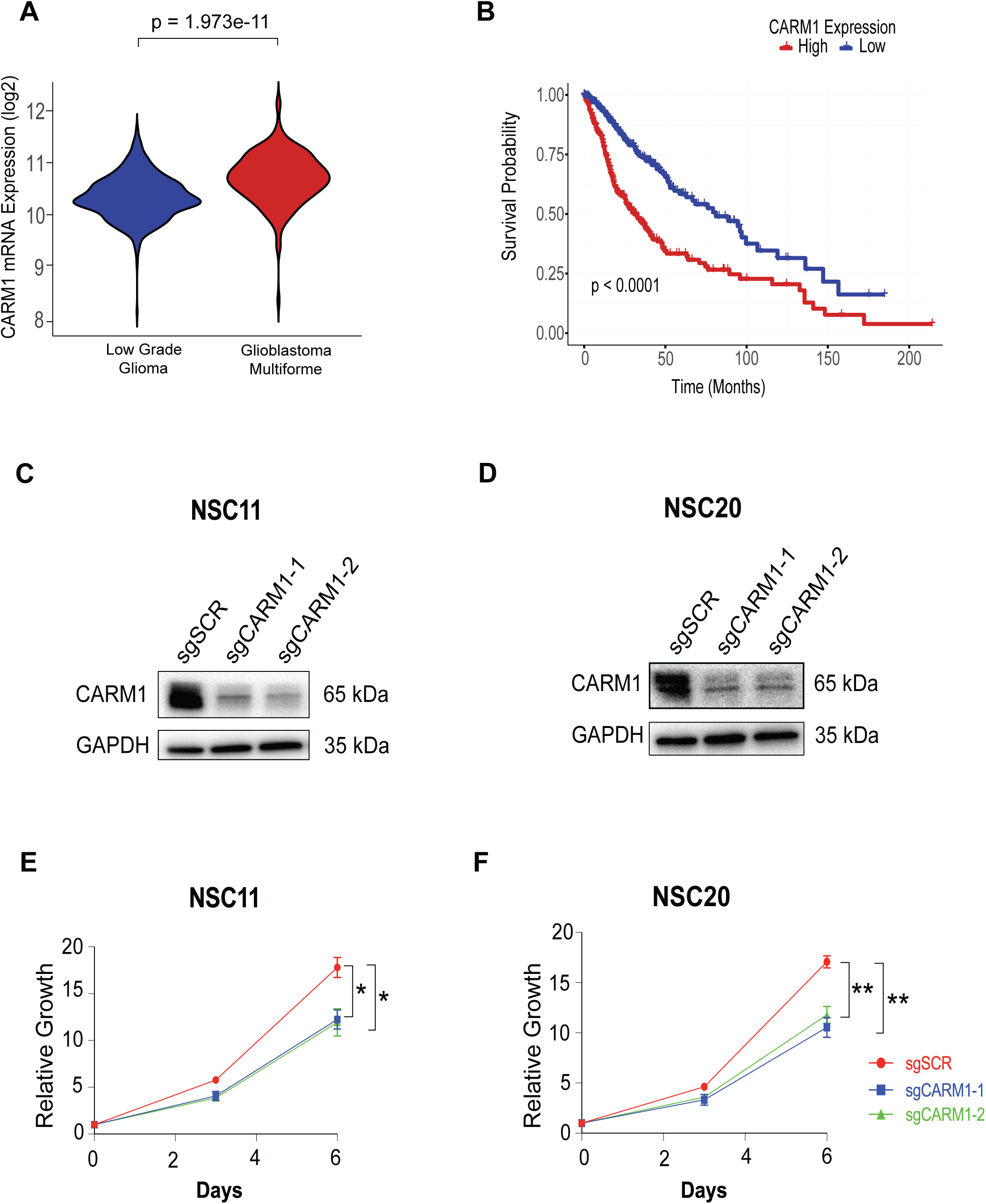
CARM1 predicts Glioma patient survival and regulates Glioma stem-like cell proliferation. **A)** Violin plot of mRNA expression log2(normalized count+1) of CARM1 in low grade glioma (n = 344) and Glioblastoma multiforme (n = 342), and **B)** Kaplan Meir curve of Glioblastoma patients with high or low CARM1 mRNA expression using UCSC Xena Browser. **C-D)** Western blot of CARM1 following CRISPR/Cas9 depletion and negative selection in NSC11 and NSC20 GSCs. **E-F)** Growth curves of sgSCR and sgCARM1 ablated NSC11 and NSC20 cells measured with Cell Titer Glo 2.0, n = 3, data shows mean +/-SEM.

### CARM1 regulates widescale transcriptome and histone changes involved in development

CARM1 has been reported to be involved in cell development and differentiation in various organ systems including the nervous system [15, 34, 35], but its role in developmental signaling in Glioma is still unknown. To identify developmental signaling programs influenced by CARM1 in GSCs we performed RNA-sequencing (RNA-seq) on NSC11 sgSCR, sgCARM1-1, and sgCARM1-2 cells. Loss of CARM1 causes significant global changes in GSC protein coding genes (adjusted p-value < .05) with a total of 1,452 differentially expressed genes (**Fig. 2A**). We find significant dysregulation of key developmental genes such as NGFR, POU3F1 (OCT6), GFAP, CHI3L1, and ARX (**Fig. 2A, supplementary table 1**). Moreover, using Gene Ontology Gene Set Enrichment Analysis we find that loss of CARM1 results in downregulation in transcriptional pathways involved in both stem cell maintenance and chromatin organization (**Fig. 2B, supplementary table 1**). In line with our results, CARM1 has been shown to be involved in histone arginine methylation, which can impact chromatin organization [36, 37]. Interestingly, loss of CARM1 results in reduced gene expression of multiple key enzymes involved in catalyzing or removing histone post-translational modifications (PTMs) besides those involved in histone arginine methylation (**Fig. 2C**). Specifically, we find that loss of CARM1 results in a reduction of key enzymes such as KDM6B, KAT2A, PHF8, and KDM2B (**Fig. 2C**) which have been shown to regulate histone H3K27me3, H3K18ac, H4K20me3, and H3K36me2 respectively. KDM6B, PHF8, and KDM2B are known histone H3/H4 lysine demethylase enzymes that remove H3K27me3, H4K20me3, and H3K36me1/2 respectively, while KAT2A is the primary catalytic lysine methyltransferase of H3K18ac [38–42]. Since these histone PTMs are often reported to be involved in cell development [43–47] and we observe strong differential expression of developmental genes upon CARM1 depletion, we hypothesized that H3K27me3, H3K18ac, and H4K20me3 would be dysregulated in sgCARM1 GSCs compared to sgSCR. To test for alterations in histone post-translational modifications we isolated histones from sgSCR and sgCARM1 cells followed by LC-MS as noted previously [23]. We observe global upregulation of co-occurring H3K27me3K36me2 and H4K20me3, and downregulation of H3K18ac (**Fig. 2D**), which is directionally in line with the differential expression observed in KDM6B, KDM2B, PHF8, and KAT2A. We also observe a profound loss in H3K79me2, a mark shown to be a transcriptional driver of developmental genes in various cell types, including heart and brain [48, 49]. We did not observe differential expression of the H3K79me2 methyltransferase, DOT1L. In summary, these results demonstrate that loss of CARM1 dysregulates the transcription of protein coding genes involved in development and chromatin organization, and creates global changes in the histone PTM architecture of GSCs.

**Figure 2:**
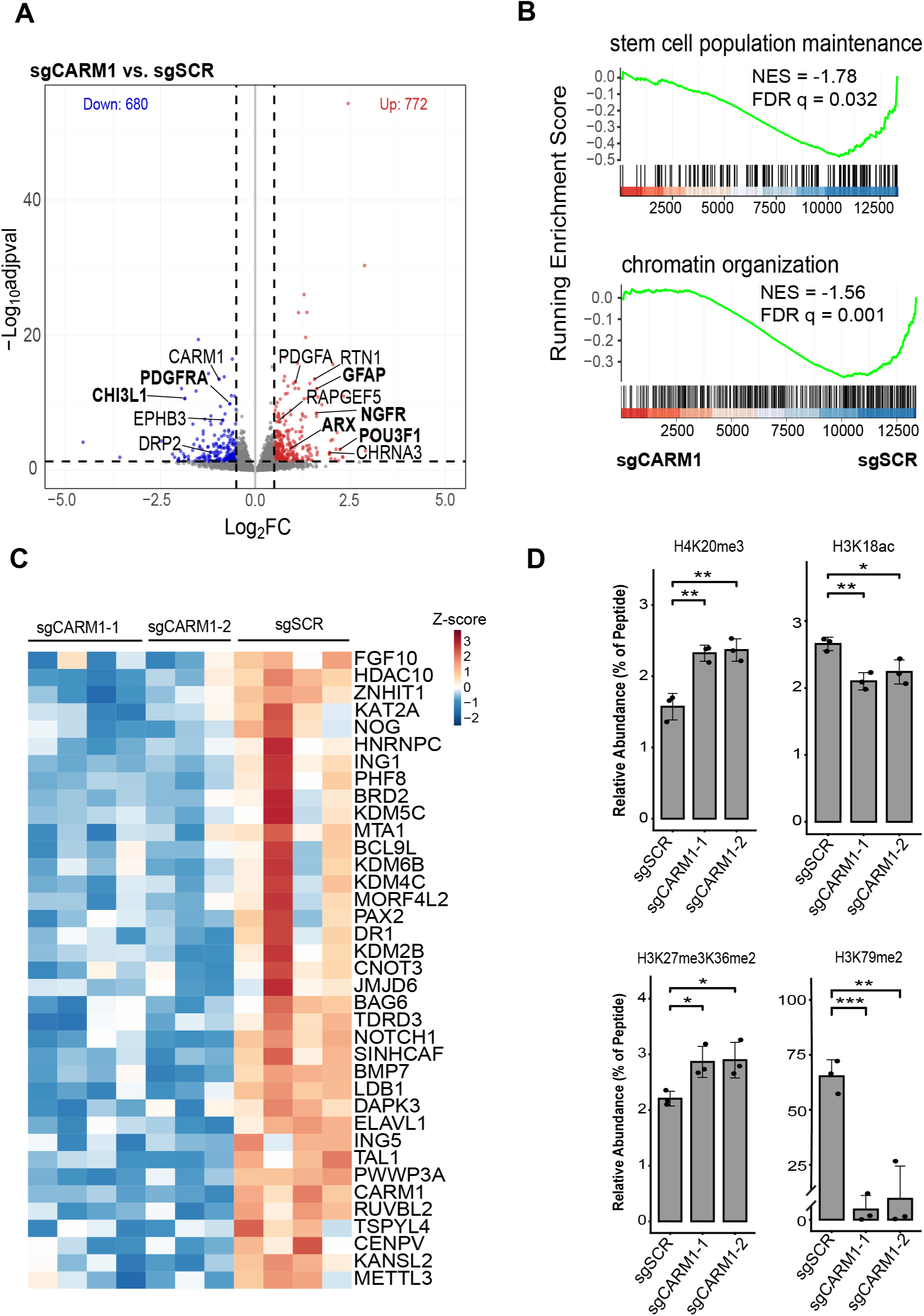
CARM1 regulates widescale transcriptome and histone PTMs involved in development. **A)** Volcano plot of NSC11 sgCARM1 (combined sgCARM1-1 and sgCARM1-2) vs sgSCR highlighting key developmental genes (adjusted p-value < 0.05, n = 3-4 per sgRNA). **B)** GSEA Enrichment plots of RNA-sequencing data of NSC11 sgSCR and sgCARM1 cells for ‘stem cell population maintenance’ (upper panel) and chromatin organization (lower panel) gene sets. **C)** Heatmap of differentially expressed genes involved in chromatin organization (adjusted p-value < 0.05) scaled by Z-score of mRNA normalized counts. **D)** Mass spectrometry analysis of significant histone post-translational modifications (PTMs) in NSC11 sgSCR and sgCARM1 cells, data shows mean +/-SD.

### CARM1 represses radial glial cell proteomic signature in Glioma stem-like cells

Since gene expression may not always correlate to protein expression, we set out to test if developmental changes were also observed on the protein level as this would give further evidence that CARM1 regulates GSC development processes. To test this, we performed global proteomics analysis via LC-MS of NSC11 sgSCR and sgCARM1 GSCs. In support of our transcriptomic findings, we identified that loss of CARM1 drives differential expression of key developmental proteins such as NGFR, GFAP, and PDGFRA, NESTIN, NOTCH1, and others (**Fig. 3A, supplementary table 2**). To test for more in depth functional protein changes we performed GSEA using C2 MSigDB. We found that loss of CARM1 drives increased expression of proteins typically enriched in human radial glial cells compared to neurons [50] (**Fig. 3B-C, supplementary table 2**), which we also observe in our transcriptomics data (**Supplementary Fig. 1A-B**). Radial glial cells are an early type of neural progenitor cells that arises during embryonic development and possess extensive migratory capabilities and astrocytic characteristics [51]. Adult human Glioma indeed maintains a population of radial glial-like cells which can promote tumor heterogeneity [52, 53]. In addition, GBM cells that adapt new cell lineages often rely on different molecular processes for growth and survival [54, 55]. To understand which molecular alterations might be the most critical to growth and survival of GSCs post CARM1 depletion, we plotted the RNA-seq and Proteomic fold changes of differentially expressed genes in both CARM1 sgRNAs. We found that NGFR was one of the most upregulated genes at both the mRNA and protein level (**Fig. 3D**). After averaging and performing rank analysis of the Proteomic and RNA log2 fold changes we observed that NGFR was the 3^rd^ most upregulated gene/protein in our data, just above GFAP—one of the known major astrocytic markers (**Fig. 3E**). NGFR has been shown to be critical in promoting migration during neurodevelopment and tumorigenesis [56, 57]. These results indicate that at both the transcriptomic and proteomic levels, CARM1 naturally represses radial glial cell signatures in GSCs and may become more dependent on NGFR signaling upon CARM1 depletion.

**Figure 3:**
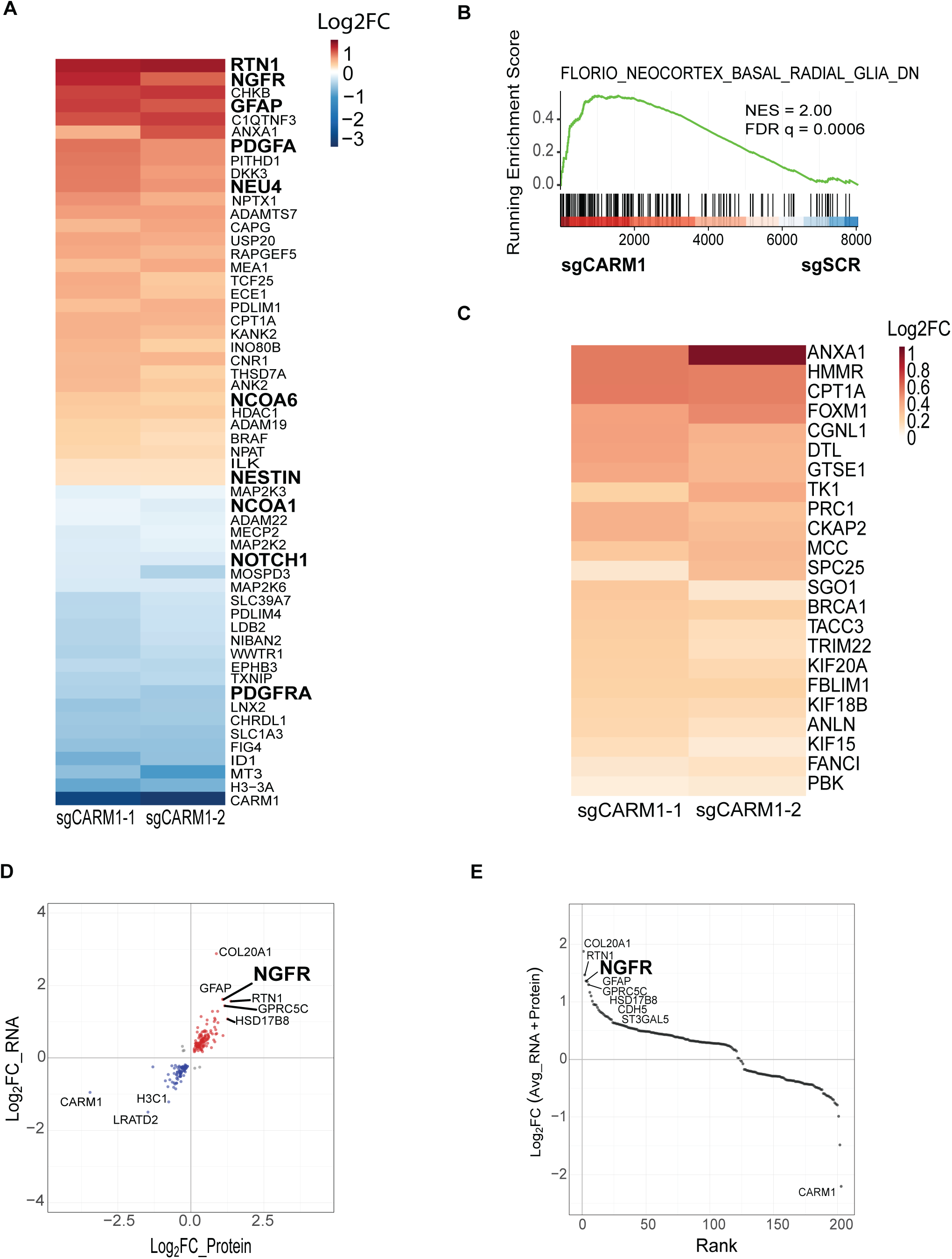
CARM1 represses radial glial cell proteomic signature in Glioma stem-like cells. **A)** Proteomics based heatmap of key differentially expressed proteins in NSC11 sgCARM1-1 and sgCARM1-2 (compared to sgSCR) scaled by log2 fold change (p-value < 0.05), n = 2 independent experiments. **B)** GSEA enrichment plot of proteomics data highlighting protein enrichment for radial glial cells in NSC11 sgCARM1 and sgSCR cells; and a **C)** heatmap of differentially expressed proteins (p-value < 0.05) that belong to the above mentioned GSEA pathway scaled by log 2 foldchange vs. control. **D)** Scatter plot of log2 fold change (averaged between sgCARM1-1 and sgCARM1-2) of RNA-seq and Proteomics data, only including genes differentially expressed in both sgCARM1-1 and sgCARM1-2 at the RNA and Protein level. **E)** Scatter plot of ranked genes/proteins based on their combined average log2 fold change between RNA and Protein.

### Loss of CARM1 drives NGFR/NTRK signaling in Glioma stem-like cells

NGFR is a transmembrane receptor for neurotrophins such as Nerve Growth Factor and stimulates activity of neurotrophin kinases (NTRK1/2/3) [58]. In addition, NTRK signaling has been shown to be critical to the survival of radial glial cells [59]. To test if loss of CARM1 leads to upregulation of NGFR/NTRK signaling in GSCs we applied mass spectrometry based phospho-proteomics. We observed that loss of CARM1 results in widespread changes in phospho-peptides (**Fig. 4A**). Using Qiagen IPA we analyzed the most enriched pathways and the predicted upstream regulators and found that NGF/NTRK signaling are both in the top enriched canonical pathways, with NGF ranking as the 2^nd^ highest predicted upstream regulator of the phospho-proteome in CARM1 depleted cells (**Fig. 4B-C, supplementary table 3**). NGFR/NTRK signaling leads directly to increased phosphorylated AKT and ERK1/2, and usually serves as the readout for increased NGFR/NTRK signaling [60]. In our data, we also see that loss of CARM1 results in significantly increased phosphorylation of proteins involved in NGFR/NTRK signaling, including AKT1 and ERK1/2 (MAPK1/3) (**Fig. 4D-E, supplementary table 3**). We then further validate that NGFR/NTRK signaling is enriched via western blot (**Fig. 4F**). These results highlight that upon loss of CARM1 GSCs increase NGFR/NTRK signaling.

**Figure 4:**
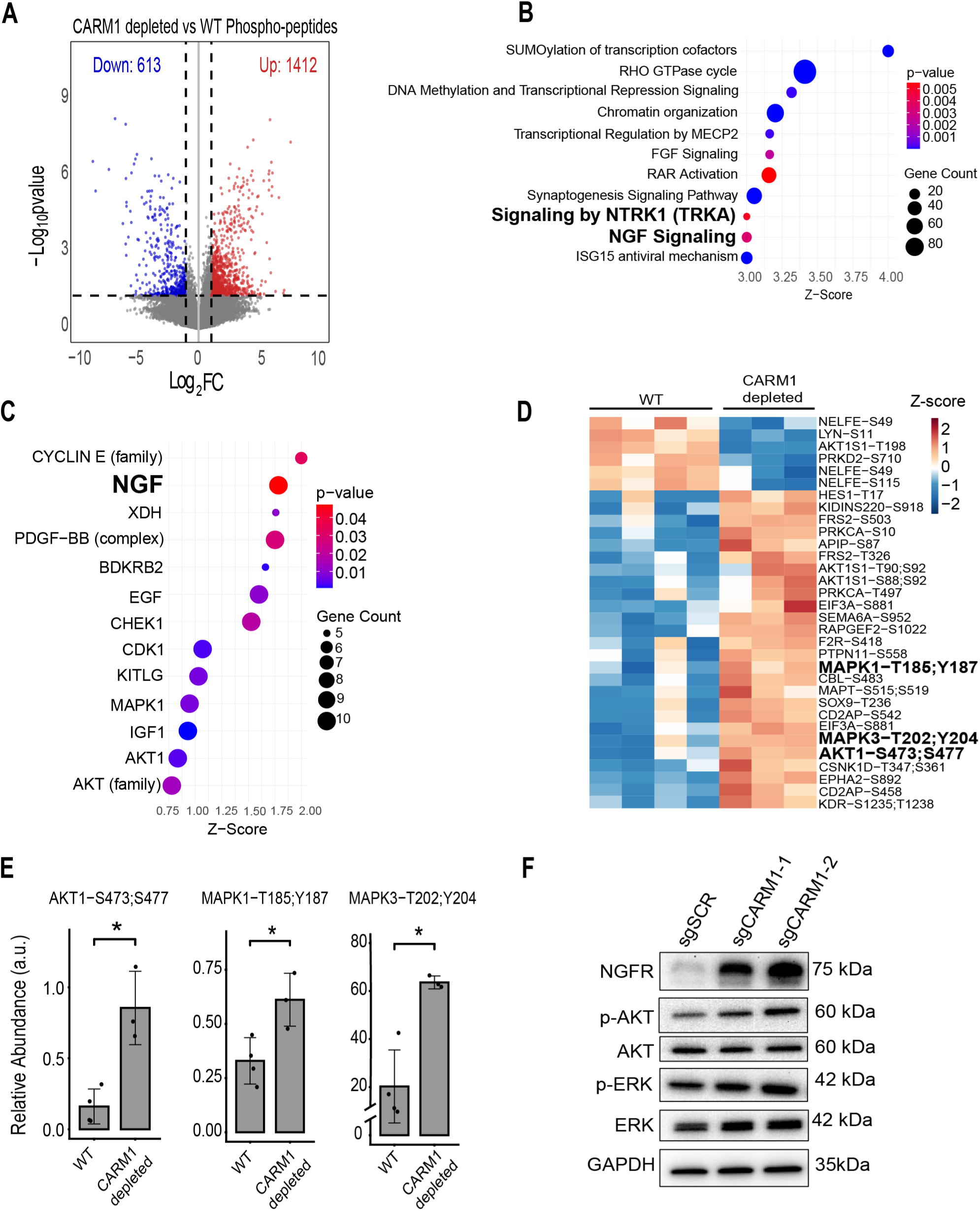
Loss of CARM1 drives NGFR/NTRK signaling in Glioma stem-like cells. **A)** Volcano plot of differentially phosphorylated peptides between NSC11 WT and CARM1 depleted cells (sgCARM1-1) with n = 3-4 (p-value < 0.05). **B)** Dot plots of Qiagen IPA analysis of top upregulated pathways **C)** and top upstream regulators ranked by Z-score. **D)** Heatmap of differentially phosphorylated proteins and phospho-sites involved in NGFR/NTRK signaling scaled by Z-score of relative abundance (p-value < 0.05) and **E)** bar plots of phosphorylation found on NGFR/NTRK targets AKT1 and MAPK1/3 (ERK1/2). **F)** Western blot validation of NGFR/NTRK signaling in NSC11 sgSCR and sgCARM1 cells.

### CARM1 substrate, NFIA, represses NGFR expression in Glioma

CARM1, like other type I PRMTs, exerts its biological effects through placing asymmetric di-methyl arginine (ADMA) on proteins and loss of ADMA has repeatedly been shown to lead to a reduction in direct protein function and/or protein-protein/protein-nucleic acid interactions [61–65]. To understand mechanistically how CARM1 might be regulating NGFR signaling we used an antibody and mass spectrometry approach to define the known CARM1 substrates in NSC11 GSCs—a method previously used to identify CARM1 substrates in human breast cancer cells [21]. Using an Asymmetric Di-Methyl Arginine (ADMA) Motif antibody we isolated peptides with ADMA in both wild-type and CARM1 depleted cells and performed LC-MS to identify CARM1 specific substrates (**Fig. 5A**). We compiled a list of known CARM1 substrates/interactors from various publications and compared our identified substrates to the available datasets. We found a total of 107 CARM1 substrates in our NSC11 GSC line, and all but two of those substrates have been shown to be arginine methylated previously [66] (**Supplementary Table 4**). Of our 107 substrates we found 56 new human CARM1 substrates (**Fig. 5B**) not identified in previous human studies on CARM1 substrates [21, 67]. To develop an understanding of the functions of the identified CARM1 substrates we performed Gene Ontology over representation analysis to identify the molecular functions. The CARM1 substrates were almost all primarily involved in the regulation of transcription (**Fig. 5C, supplementary table 4**). We find that NFIA and NFIX are two of the most enriched CARM1 substrates in our cell line (**Fig. 5D**). NFIA and NFIX are transcription factors that regulate neural development [68, 69] and have been confirmed to be CARM1 substrates in mouse embryonic fibroblast [70]. Moreover, using ChIP-seq it has previously been shown that NGFR is a target of NFIA and NFIX [71]. Since NFIA specifically has been shown to be significantly involved in Glioma tumorigenesis [72–75] we hypothesized NFIA to be the primary CARM1 substrate responsible for regulating NGFR signaling in GSCs. To test if NFIA is able to regulate NGFR expression in GSCs we used siRNA to knockdown NFIA and probed NGFR expression with western blot. We find that knockdown of NFIA resulted in an upregulation of NGFR expression (**Fig. 5E**), analogous to CARM1 depletion (**Fig. 5F**). These results show that NFIA is a CARM1 substrate in GSCs and represses NGFR signaling just as CARM1 does, therefore suggesting that NFIA is the likely CARM1 substrate responsible for NGFR regulation in GSCs.

**Figure 5:**
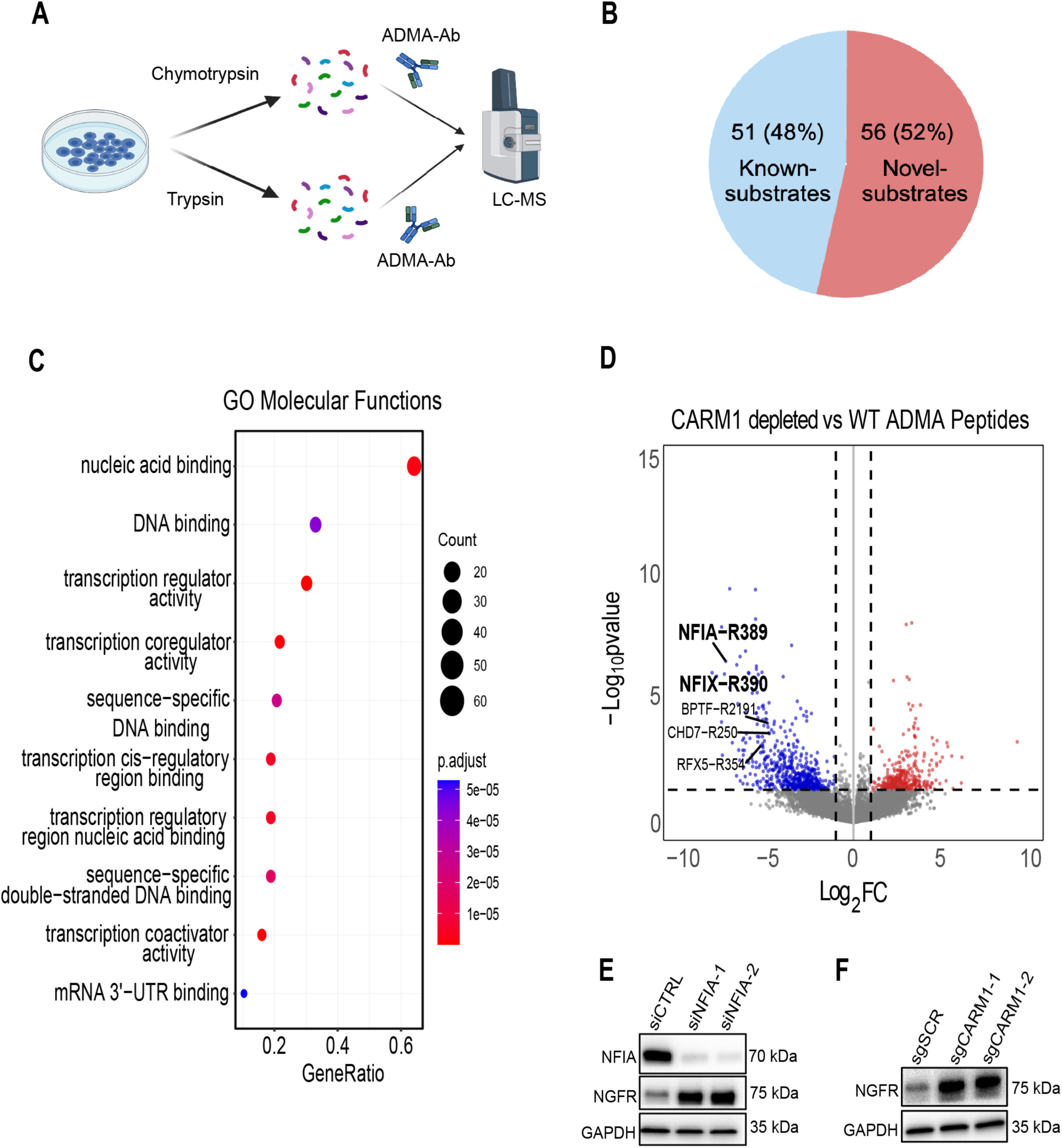
CARM1 substrate, NFIA, represses NGFR expression. **A)** Mass spectrometry schematic of approach to identify CARM1 asymmetric di-methyl arginine (ADMA) substrates in NSC11 GSCs WT vs CARM1-depleted (sgCARM1-1), n = 3-4. **B)** Pie chart of identified CARM1 substrates. **C)** Dot plot of Gene Ontology Molecular Functions of statistically significant (p-value < 0.05) CARM1 substrates and **D)** volcano plot of differentially ADMA peptides, labeling key substrates (i.e. NFIA). **E)** Western blots of NFIA and NGFR expression after knockdown of NFIA and **F)** of NGFR expression post CARM1 depletion.

### Loss of CARM1 sensitizes GSCs to NTRK inhibition

Our work so far has shown that CARM1 has a natural repressive effect on NGFR/NTRK signaling in GSCs, so we hypothesized that upon NTRK inhibition GSCs would downregulate CARM1 as a compensatory mechanism to maintain NGFR/NTRK homeostasis. To test this, we treated NSC11 GSCs with DMSO or the NTRK inhibitor Entrectinib for 48 hours. As proposed, GSCs downregulate CARM1 upon NTRK inhibition, likely to compensate for the reduction in NTRK inhibition (**Fig. 6A**). Since loss of CARM1 drives NGFR/NTRK signaling in GSCs and NGFR/NTRK promote differentiation and survival [60], we hypothesized that GSCs lacking CARM1 would have increased sensitivity to NTRK inhibition. Currently there are several NTRK inhibitors that exist, but the compound Entrectinib is both FDA approved and brain-penetrant [76–78], so we use Entrectinib as our primary treatment modality. To test if CARM1 loss leads to increased short-term sensitivity of GSCs to NTRK inhibition we treated GSCs with Entrectinib for 96 hours and measured cell viability. We found that sgCARM1 GSCs have increased sensitivity to Entrectinib compared to sgSCR (**Fig. 6B**). We also validate that Entrectinib sensitivity is dependent on NGFR expression by demonstrating that GSCs with siRNA knockdown of NGFR have increased resistance to Entrectinib (**Supplementary Fig. 2**). Since AKT is also activated by NGFR/NTRK signaling we also treated GSCs with the FDA approved AKT inhibitor Capivasertib and saw that sgCARM1 GSCs had increased sensitivity to AKT inhibition compared to sgSCR (**Supplementary Fig. 3).** We next checked long term proliferative capacity by treating GSCs with 3uM of Entrectinib (ic50 of sgCARM1 cells) using the clonogenic assay. We found that sgCARM1 cells have a reduced baseline colony formation compared to sgSCR under DMSO conditions (**Fig. 6C)**, in line with our earlier results demonstrating sgCARM1 GSCs have reduced proliferation. We also discovered that sgCARM1 GSCs are over 4 fold more sensitive to Entrectinib compared to sgSCR cells (**Fig. 6D**). These results demonstrate that loss of CARM1 in GSCs creates an increased sensitivity to Entrectinib in both the short and long term in-vitro.

**Figure 6:**
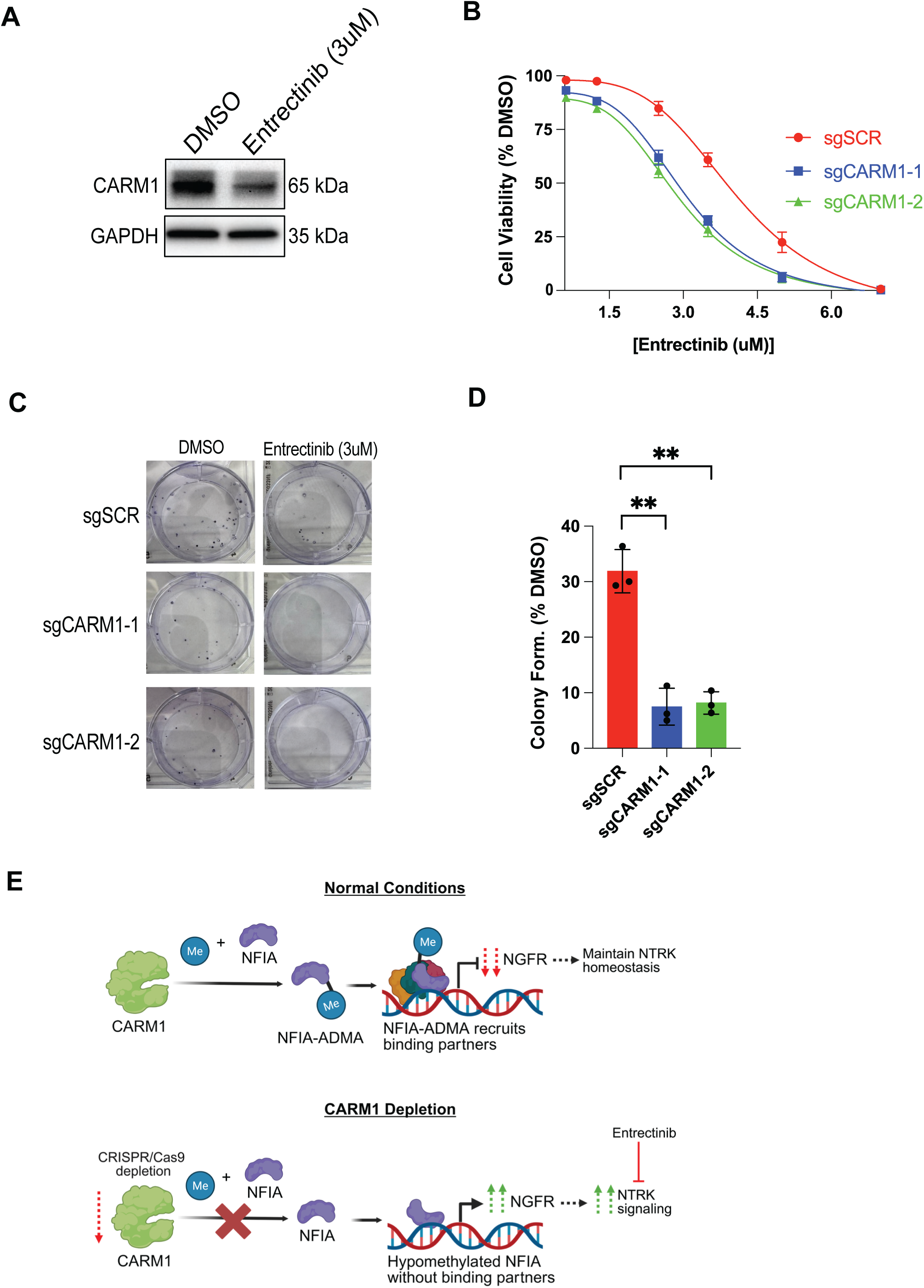
Loss of CARM1 sensitizes Glioma stem-like cells to NTRK inhibition. **A)** Western blot of CARM1 expression after 48 hours treatment with DMSO or Entrectinib (3uM). **B)** Cell viability of NSC11 GSCs after 96 hours of Entrectinib measured with Cell Titer Glo 2.0, n = 3, data shows mean +/- SEM. **C-D)** Clonogenic assay of NSC11 GSCs treated for 72 hours with Entrectinib, n = 3, data shows mean +/- SD. **E)** *(upper panel)* Under normal conditions we propose CARM1 methylates NFIA on R389, making NFIA-ADMA. NFIA-ADMA is then able to properly form various protein-protein interactions, possibly with co-repressors, to repress NGFR transcription. Reduced NGFR transcription maintains homeostatic levels of NTRK signaling. *(lower panel)* Upon CARM1 depletion, NFIA is not methylated and fails to form proper protein-protein interactions. Therefore, when hypomethylated-NFIA binds chromatin, it is unable to successfully repress NGFR. NGFR expression increases which then promotes NTRK signaling which can be targeted with the NTRK inhibitor, Entrectinib.

## Discussion

Our work set out to discover if CARM1 is a reasonable therapeutic target for Glioblastoma and how CARM1 regulates GSC character and developmental processes. We found that CARM1 has a positive regulation on GSC cell proliferation and its mRNA expression strongly correlates with GBM patient survival, with lower CARM1 expression predicting better survival. We discovered that CARM1 also had a profound impact on the gene expression of developmental and chromatin organizing genes, and regulates the histone PTM profile of GSCs beyond the known H3R17 methylation. Using both Proteomics and RNA-seq we reveal that CARM1 prevents GSCs from shifting to the radial glial cell lineage, altering their fate commitment. With phospho-proteomics we found that depletion of CARM1 results in strong upregulation of NGFR/NTRK signaling. When CARM1 is depleted from GSCs they not only upregulate NGFR/NTRK but they also become highly dependent on this signaling cascade for cell survival, evidenced by sgCARM1 cells increased sensitivity to NTRK and AKT inhibitors. Mechanistically, using mass spectrometry we reveal that NFIA is one of the most regulated CARM1 methylation substrates in our GSCs, and that knockdown of NFIA recapitulates NGFR overexpression observed in sgCARM1 cells. This suggest that the CARM1-NFIA relationship naturally represses NGFR signaling and when lost, results in upregulation of NGFR signaling in GSCs.

Though CARM1 has been investigated largely for its role in normal tissue development, less focus has been placed on its use as a therapeutic target. One issue with targeting PRMTs is that there is redundancy in their substrates, and so when one PRMT is inhibited others can compensate. This is not the case for CARM1. CARM1 methylates proline-arginine rich motifs, while other PRMTs methylate glycine-arginine rich motifs thus CARM1’s differences in motif preference makes its lost difficult to compensate [14, 79]. Moreover, CARM1 depletion has significantly less toxicity in-vivo compared to depletion of other PRMTs (i.e. PRMT1 or PRMT5) and thus may be more tolerable in the clinic [14, 80, 81]. Our work demonstrates that loss of CARM1 causes a reduction in GSC cell proliferation and gives support to the proposal that CARM1 should be further investigated as a therapeutic target in Glioblastoma, perhaps focusing on the stem-like population.

CARM1 has been previously reported to affect chromatin organization through its methylation of H3R17 and/or H3R26, but our work indeed reveals that CARM1 regulates several other histone PTMs as well. We observe that CARM1 depletion leads to an upregulation of H4K20me3 and bivalent H3K27me3K36me2, coupled with a downregulation of H3K18ac and H3K79me2. Likewise, we found differential expression of key enzymes that regulate these histone PTMs (i.e. PHF8, KDM6B, KDM2B, KAT2A). Thus, CARM1 appears to regulate more histone PTMs than previously thought, and it likely does this through transcriptional control of various histone PTM readers/writers/erasers. These histone PTMs have been shown to play various roles in cellular development. H4K20me3 is often found on heterochromatin and associated with cell lineage commitment in embryonic stem cells [82]. H3K27me3K36me2 is a bivalent histone mark that has been observed to be enriched in mesenchymal stem cells [83]. Though this mark includes H3K36me3, our work has previously shown via mass spectrometry based histone analysis and Ch-IP/RNA-seq that bivalent H3K27me3K36me2 has a dominant repressive effect on transcription and chromatin accessibility [84]. H3K18ac and H3K79me2 are histone PTMs often found at the TSS of genes undergoing active transcription and have meaningful roles in cellular development [48, 85, 86]. The shifts in the histone PTM architecture upon CARM1 depletion are evident of a changing chromatin landscape in which there is a gain of repressive/heterochromatin marks and a loss of transcriptional activation/euchromatin marks. This is line with the work of Mark Bedford’s group showing that CARM1 can promote an open chromatin state in small cell lung cancer models [70]. If indeed CARM1 depletion leads to a reduction in chromatin accessibility via histone PTM remodeling this may have strong implications on the stem-like nature of GSCs, their lineage potential, and therapy resistance.

At both the transcript and protein level we observe upregulation of various developmental markers like RTN1 and GFAP, and downregulation of markers such as NOTCH1 and PDGFRA. To our surprise we found an increase in genes/proteins enriched in the radial glial cell lineage upon CARM1 depletion. This suggest that in GSCs loss of CARM1 does not necessary lead to complete cell differentiation but instead leads to a shift in stem-like cell lineage promoting a more radial glial like cell type in Glioma. Radial glial cells have been discovered in Gliomas and play a key role in establishing intra-tumoral heterogeneity [53]. The shift to a more radial glial-like cell type may not be optimal for glioma therapy as this cell type is more invasive [53]. However, though radial glial cells are more invasive it is not clear that they contribute to worse patient outcomes, as Glioblastoma is already highly invasive by nature and is typically spread throughout the brain at diagnosis. Despite the gaps in knowledge that exist about radial glial cells in Glioblastoma, to our knowledge, we are the first group to reveal that CARM1 regulates the transcriptomic/proteomic radial glial lineage in GSCs and may shed new light on the fundamental characteristics of GSC plasticity.

One of the most upregulated genes/proteins upon CARM1 depletion in GSCs was NGFR. NGFR is a transmembrane receptor that stimulates the activity of NTRK1/2/3 and is vital to cell migration during neurodevelopment and tumorigenesis [56–58]. Moreover, phospho-proteomics and western blot revealed that sgCARM1 cells have upregulation of the entire NGFR/NTRK pathway. NTRK signaling plays a vital role in promoting the regeneration and survival of radial glial cells [59], thus in line with our results showing that CARM1 depleted GSCs shift their lineage towards radial glial cells. This newfound dependency on NGFR/NTRK signaling upon CARM1 loss can be exploited in-vitro with direct inhibition of NTRK via Entrectinib or through downstream inhibition of AKT kinase with Capivasertib. This work highlights the need to develop a deeper understanding of the cell lineage regulators in GSCs as they may unlock new therapeutic strategies in Glioblastoma. Although there are single cell technologies currently being used to help provide insight into lineage specific alterations in Glioblastoma, these studies often don’t validate the dependencies observed in each lineage or the genetic/epigenetic regulators of each lineage. Developing an understanding of which genetic/epigenetic regulators allow GSCs to switch between various neural lineages may help personalize our therapeutics and better predict reoccurrence. Further, to increase clinical relevance, more studies should be performed that target various cell lineages within Glioblastoma with chemical or CRISPR/Cas9 screens to validate which pathways are not just differentially expressed but necessary for survival.

Our work along with others have shown that NFIA is a CARM1 substrate [70], and in our case it is one of the most differentially methylated CARM1 substrates in our dataset. Loss of protein arginine methylation has repeatedly been shown to disrupt protein function and cellular processes [62, 87–89], and we expected that the profound loss of NFIA R389 methylation (log2FC < -7 in CARM1 depleted vs WT) would lead to a disruption of NFIA’s transcriptional functions. Further, the fact that NGFR promoter is bound by NFIA [71] made us suspect that it was the likely CARM1 substrate responsible for regulating NGFR. Knockdown of NFIA indeed recapitulated our observation that CARM1 depletion drives up NGFR expression. Without a mutation of R389 on NFIA we cannot definitively confirm that the loss of methylation of NFIA is the driving cause of NGFR upregulation. However, we do show that the CARM1/NFIA relationship is the likely upstream regulator of NGFR in GSCs. More studies should be performed to verify if the methylation on NFIA R389 itself is the cause of NGFR dysregulation upon CARM1 loss and how that loss of methylation affects protein function. Based on alpha fold predictions R389 is not on the DNA binding region of NFIA and we don’t find that NFIA is reduced upon chromatin after CARM1 loss (data not shown). So, we suspect that this loss of methylation may impact NFIA protein interactions (**Fig. 6E**). It is possible that the loss of R389 methylation disrupts NFIA ability to bind co-repressors, allowing NGFR expression to increase (**Fig. 6E**).

In conclusion, our results reveal that CARM1 regulates both the gene and protein expression of key developmental markers in GSCs and specifically represses GSCs from shifting to the radial glial cell lineage. The NGFR/NTRK signaling cascade, a signaling pathway vital to radial glial cell survival, becomes upregulated upon CARM1 loss. Moreover, the upregulation of NGFR/NTRK signaling upon CARM1 loss is likely due to profound loss of CARM1 dependent methylation of R389 on NFIA. Finally, we then show that not only does CARM1 loss upregulate NGFR/NTRK signaling but also that CARM1 loss sensitizes GSCs to NTRK (Entrectinib) and AKT (Capivasertib) inhibition in-vitro. Ultimately this work sheds new light on how GSCs regulate their cellular fate and how these fundamental mechanisms can be targeted therapeutically.

## Raw data availability

The mass spectrometry data have been deposited to the ProteomeXchange Consortium via the PRIDE [90] partner repository. Raw sequencing data has been deposited in the NCBI Sequence Read Archive under BioProject accession PRJNA1237514. Processed gene expression data has been deposited in the NCBI Gene Expression Omnibus database under accession GSE292789.

## Acknowledgments

The Sidoli lab gratefully acknowledges for funding the Hevolution Foundation (AFAR), the Einstein-Mount Sinai Diabetes center, the NIH Office of the Director (S10OD030286), and the NIH SPORE (P50CA127001) in supplying the GSC cell lines.

## Authors’ Contributions

Conception and design: D.L. Young, C. Guha, S. Sidoli

Development of methodology: D.L. Young, R.E. Phillips, D. Shechter, S. Sidoli, C. Guha, J. Aguilan, P.J. Tofilon

Acquisition of data: D.L. Young, J. Roth, J. DeAngelo, B. Bell, J. Vercellino, S. Stransky, B. Malachowska

Analysis and interpretation: D.L. Young., R. Cutler., S. Sidoli, C. Guha, R.E. Phillips

Writing and revision of manuscript: D.L. Young, R. Cutler, B. Malachowska, J. Vercellino, S. Sidoli, C. Guha

Study supervision: S. Sidoli, C. Guha

## References

1. Poon, M.T.C., et al., Longer-term (>/= 2 years) survival in patients with glioblastoma in population-based studies pre- and post-2005: a systematic review and meta-analysis. Sci Rep, 2020. 10(1): p. 11622.

2. Weller, M., et al., Standards of care for treatment of recurrent glioblastoma--are we there yet? Neuro Oncol, 2013. 15(1): p. 4–27.

3. Prager, B.C., et al., Glioblastoma Stem Cells: Driving Resilience through Chaos. Trends Cancer, 2020. 6(3): p. 223–235.

4. Singh, S.K., et al., Identification of human brain tumour initiating cells. Nature, 2004. 432(7015): p. 396–401.

5. Valor, L.M. and I. Hervás-Corpión, The Epigenetics of Glioma Stem Cells: A Brief Overview. Frontiers in Oncology, 2020. 10.

6. Phillips, R.E., A.A. Soshnev, and C.D. Allis, Epigenomic Reprogramming as a Driver of Malignant Glioma. Cancer Cell, 2020. 38(5): p. 647–660.

7. Liau, B.B., et al., Adaptive Chromatin Remodeling Drives Glioblastoma Stem Cell Plasticity and Drug Tolerance. Cell Stem Cell, 2017. 20(2): p. 233–246 e7.

8. Bryant, J.-P., J. Heiss, and Y.K. Banasavadi-Siddegowda, Arginine Methylation in Brain Tumors: Tumor Biology and Therapeutic Strategies. Cells, 2021. 10(1): p. 124.

9. Blanc, R.S. and S. Richard, Arginine Methylation: The Coming of Age. Molecular Cell, 2017. 65(1): p. 8–24.

10. Guccione, E. and S. Richard, The regulation, functions and clinical relevance of arginine methylation. Nature Reviews Molecular Cell Biology, 2019. 20(10): p. 642–657.

11. Banasavadi-Siddegowda, Y.K., et al., PRMT5–PTEN molecular pathway regulates senescence and self-renewal of primary glioblastoma neurosphere cells. Oncogene, 2017. 36(2): p. 263–274.

12. Zheng, D., et al., LncRNA NNT-AS1 promote glioma cell proliferation and metastases through miR-494-3p/PRMT1 axis. Cell Cycle, 2020. 19(13): p. 1621–1631.

13. Sachamitr, P., et al., PRMT5 inhibition disrupts splicing and stemness in glioblastoma. Nature Communications, 2021. 12(1).

14. Santos, M., J.W. Hwang, and M.T. Bedford, CARM1 arginine methyltransferase as a therapeutic target for cancer. Journal of Biological Chemistry, 2023. 299(9): p. 105124.

15. Wu, Q., et al., CARM1 is Required in Embryonic Stem Cells to Maintain Pluripotency and Resist Differentiation. Stem Cells, 2009. 27(11): p. 2637–2645.

16. Torres-Padilla, M.-E., et al., Histone arginine methylation regulates pluripotency in the early mouse embryo. Nature, 2007. 445(7124): p. 214–218.

17. Rios, A.F.L., et al., Expression of pluripotency-related genes in human glioblastoma. Neurooncol Adv, 2022. 4(1): p. p.

18. McCord, A.M., et al., CD133+ glioblastoma stem-like cells are radiosensitive with a defective DNA damage response compared with established cell lines. Clin Cancer Res, 2009. 15(16): p. 5145–53.

19. Ran, F.A., et al., Genome engineering using the CRISPR-Cas9 system. Nature Protocols, 2013. 8(11): p. 2281–2308.

20. Franken, N.A.P., et al., Clonogenic assay of cells in vitro. Nature Protocols, 2006. 1(5): p. 2315–2319.

21. Shishkova, E., et al., Global mapping of CARM1 substrates defines enzyme specificity and substrate recognition. Nature Communications, 2017. 8(1): p. 15571.

22. Mayoral, J., et al., Toxoplasma gondii PPM3C, a secreted protein phosphatase, affects parasitophorous vacuole effector export. PLOS Pathogens, 2020. 16(12): p. e1008771.

23. Joseph-Chowdhury, J.N., et al., Global Level Quantification of Histone Post-Translational Modifications in a 3D Cell Culture Model of Hepatic Tissue. J Vis Exp, 2022(183).

24. Stransky, S., et al., Investigation of reversible histone acetylation and dynamics in gene expression regulation using 3D liver spheroid model. Epigenetics & Chromatin, 2022. 15(1).

25. Yu, G., et al., clusterProfiler: an R Package for Comparing Biological Themes Among Gene Clusters. OMICS: A Journal of Integrative Biology, 2012. 16(5): p. 284–287.

26. Goldman, M.J., et al., Visualizing and interpreting cancer genomics data via the Xena platform. Nature Biotechnology, 2020. 38(6): p. 675–678.

27. Demichev, V., et al., dia-PASEF data analysis using FragPipe and DIA-NN for deep proteomics of low sample amounts. Nature Communications, 2022. 13(1).

28. Perez-Riverol, Y., et al., The PRIDE database resources in 2022: a hub for mass spectrometry-based proteomics evidences. Nucleic Acids Research, 2022. 50(D1): p. D543–D552.

29. Yuan, Z.F., et al., EpiProfile 2.0: A Computational Platform for Processing Epi-Proteomics Mass Spectrometry Data. J Proteome Res, 2018. 17(7): p. 2533–2541.

30. Ewels, P.A., et al., The nf-core framework for community-curated bioinformatics pipelines. Nature Biotechnology, 2020. 38(3): p. 276–278.

31. Hart, T., et al., Finding the active genes in deep RNA-seq gene expression studies. BMC Genomics, 2013. 14(1): p. 778.

32. Krämer, A., et al., Causal analysis approaches in Ingenuity Pathway Analysis. Bioinformatics, 2014. 30(4): p. 523–530.

33. Wang, F., et al., WDR5-Myc axis promotes the progression of glioblastoma and neuroblastoma by transcriptional activating CARM1. Biochem Biophys Res Commun, 2020. 523(3): p. 699–706.

34. Ishino, Y., et al., Coactivator-associated arginine methyltransferase 1 controls oligodendrocyte differentiation in the corpus callosum during early brain development. Developmental Neurobiology, 2022. 82(3): p. 245–260.

35. Lim, C.S. and D.L. Alkon, Inhibition of coactivator-associated arginine methyltransferase 1 modulates dendritic arborization and spine maturation of cultured hippocampal neurons. J Biol Chem, 2017. 292(15): p. 6402–6413.

36. Bauer, U.M., et al., Methylation at arginine 17 of histone H3 is linked to gene activation. EMBO reports, 2002. 3(1): p. 39–44.

37. Wu, J., et al., A Role for CARM1-Mediated Histone H3 Arginine Methylation in Protecting Histone Acetylation by Releasing Corepressors from Chromatin. PLoS ONE, 2012. 7(6): p. e34692.

38. Park, S., et al., Histone lysine methylation modifiers controlled by protein stability. Experimental & Molecular Medicine, 2024. 56(10): p. 2127–2144.

39. Chung, N., et al., Active H3K27me3 demethylation by KDM6B is required for normal development of bovine preimplantation embryos. Epigenetics, 2017. 12(12): p. 1048–1056.

40. Hyun, K., et al., Writing, erasing and reading histone lysine methylations. Exp Mol Med, 2017. 49(4): p. e324.

41. Kuo, Y.-M. and A.J. Andrews, Quantitating the Specificity and Selectivity of Gcn5-Mediated Acetylation of Histone H3. PLoS ONE, 2013. 8(2): p. e54896.

42. Tsukada, Y.-I., et al., Histone demethylation by a family of JmjC domain-containing proteins. Nature, 2006. 439(7078): p. 811–816.

43. Buontempo, S., et al., EZH2-Mediated H3K27me3 Targets Transcriptional Circuits of Neuronal Differentiation. Frontiers in Neuroscience, 2022. 16.

44. Vezzoli, M., et al., TFIIIC as a Potential Epigenetic Modulator of Histone Acetylation in Human Stem Cells. International Journal of Molecular Sciences, 2023. 24(4): p. 3624.

45. Funk, O.H., et al., Postmitotic accumulation of histone variant H3.3 in new cortical neurons establishes neuronal chromatin, transcriptome, and identity. Proceedings of the National Academy of Sciences, 2022. 119(32).

46. Rajagopalan, K.N., et al., Depletion of H3K36me2 recapitulates epigenomic and phenotypic changes induced by the H3.3K36M oncohistone mutation. Proceedings of the National Academy of Sciences, 2021. 118(9): p. e2021795118.

47. Zaghi, M., V. Broccoli, and A. Sessa, H3K36 Methylation in Neural Development and Associated Diseases. Frontiers in Genetics, 2020. 10.

48. Cattaneo, P., et al., DOT1L-mediated H3K79me2 modification critically regulates gene expression during cardiomyocyte differentiation. Cell Death & Differentiation, 2016. 23(4): p. 555–564.

49. Ferrari, F., et al., DOT1L-mediated murine neuronal differentiation associates with H3K79me2 accumulation and preserves SOX2-enhancer accessibility. Nature Communications, 2020. 11(1).

50. Florio, M., et al., Human-specific gene ARHGAP11B promotes basal progenitor amplification and neocortex expansion. Science, 2015. 347(6229): p. 1465–1470.

51. Miranda-Negrón, Y. and J.E. García-Arrarás, Radial glia and radial glia-like cells: Their role in neurogenesis and regeneration. Frontiers in Neuroscience, 2022. 16.

52. Wang, R., et al., Adult Human Glioblastomas Harbor Radial Glia-like Cells. Stem Cell Reports, 2020. 14(2): p. 338–350.

53. Bhaduri, A., et al., Outer Radial Glia-like Cancer Stem Cells Contribute to Heterogeneity of Glioblastoma. Cell Stem Cell, 2020. 26(1): p. 48–63 e6.

54. Lu, X., et al., Cell-lineage controlled epigenetic regulation in glioblastoma stem cells determines functionally distinct subgroups and predicts patient survival. Nature Communications, 2022. 13(1).

55. Alcantara Llaguno, S.R., et al., Adult Lineage-Restricted CNS Progenitors Specify Distinct Glioblastoma Subtypes. Cancer Cell, 2015. 28(4): p. 429–440.

56. Bentley, C.A. and K.-F. Lee, p75 Is Important for Axon Growth and Schwann Cell Migration during Development. The Journal of Neuroscience, 2000. 20(20): p. 7706–7715.

57. Boiko, A.D., et al., Human melanoma-initiating cells express neural crest nerve growth factor receptor CD271. Nature, 2010. 466(7302): p. 133–137.

58. Barker, P.A., p75NTR is positively promiscuous: novel partners and new insights. Neuron, 2004. 42(4): p. 529–33.

59. Harada, C., et al., Glia- and neuron-specific functions of TrkB signalling during retinal degeneration and regeneration. Nature Communications, 2011. 2(1): p. 189.

60. Cocco, E., M. Scaltriti, and A. Drilon, NTRK fusion-positive cancers and TRK inhibitor therapy. Nature Reviews Clinical Oncology, 2018. 15(12): p. 731–747.

61. Veazey, K.J., et al., CARM1 inhibition reduces histone acetyltransferase activity causing synthetic lethality in CREBBP/EP300-mutated lymphomas. Leukemia, 2020. 34(12): p. 3269–3285.

62. Pham, H.Q.H., X. Tao, and Y. Yang, Protein arginine methylation in transcription and epigenetic regulation. Frontiers in Epigenetics and Epigenomics, 2023. 1.

63. Yang, Y., et al., Arginine methylation facilitates the recruitment of TOP3B to chromatin to prevent R loop accumulation. Mol Cell, 2014. 53(3): p. 484–97.

64. Nie, M., et al., CARM1-mediated methylation of protein arginine methyltransferase 5 represses human γ-globin gene expression in erythroleukemia cells. Journal of Biological Chemistry, 2018. 293(45): p. 17454–17463.

65. Lorton, B.M., et al., A Binary Arginine Methylation Switch on Histone H3 Arginine 2 Regulates Its Interaction with WDR5. Biochemistry, 2020. 59(39): p. 3696–3708.

66. Maron, M.I., et al., Independent transcriptomic and proteomic regulation by type I and II protein arginine methyltransferases. iScience, 2021. 24(9): p. 102971.

67. Itonaga, H., et al., Tyrosine phosphorylation of CARM1 promotes its enzymatic activity and alters its target specificity. Nature Communications, 2024. 15(1).

68. Heng, Y.H.E., et al., NFIX Regulates Neural Progenitor Cell Differentiation During Hippocampal Morphogenesis. Cerebral Cortex, 2014. 24(1): p. 261–279.

69. Piper, M., et al., NFIA Controls Telencephalic Progenitor Cell Differentiation through Repression of the Notch Effector Hes1. Journal of Neuroscience, 2010. 30(27): p. 9127–9139.

70. Gao, G., et al., The NFIB/CARM1 partnership is a driver in preclinical models of small cell lung cancer. Nature Communications, 2023. 14(1).

71. Fraser, J., et al., Common Regulatory Targets of NFIA, NFIX and NFIB during Postnatal Cerebellar Development. The Cerebellum, 2020. 19(1): p. 89–101.

72. Chen, K.S., et al., NFIA and NFIB function as tumour suppressors in high-grade glioma in mice. Carcinogenesis, 2021. 42(3): p. 357–368.

73. Glasgow, S.M., et al., Glia-specific enhancers and chromatin structure regulate NFIA expression and glioma tumorigenesis. Nature Neuroscience, 2017. 20(11): p. 1520–1528.

74. Lee, J., E. Hoxha, and H.-R. Song, A novel NFIA-NFκB feed-forward loop contributes to glioblastoma cell survival. Neuro-Oncology, 2016: p. now233.

75. Glasgow, S.M., et al., Mutual antagonism between Sox10 and NFIA regulates diversification of glial lineages and glioma subtypes. Nature Neuroscience, 2014. 17(10): p. 1322–1329.

76. Grogan, P.T., et al., Entrectinib demonstrates prolonged efficacy in an adult case of radiation-refractory NTRK fusion glioblastoma. Neuro-Oncology Advances, 2022. 4(1).

77. Fischer, H., et al., Entrectinib, a TRK/ROS1 inhibitor with anti-CNS tumor activity: differentiation from other inhibitors in its class due to weak interaction with P-glycoprotein. Neuro-Oncology, 2020. 22(6): p. 819–829.

78. Desai, A.V., et al., Entrectinib in children and young adults with solid or primary CNS tumors harboring NTRK, ROS1, or ALK aberrations (STARTRK-NG). Neuro-Oncology, 2022. 24(10): p. 1776–1789.

79. Cheng, D., et al., The Arginine Methyltransferase CARM1 Regulates the Coupling of Transcription and mRNA Processing. Molecular Cell, 2007. 25(1): p. 71–83.

80. Yu, Z., et al., A Mouse PRMT1 Null Allele Defines an Essential Role for Arginine Methylation in Genome Maintenance and Cell Proliferation. Molecular and Cellular Biology, 2009. 29(11): p. 2982–2996.

81. Tee, W.W., et al., Prmt5 is essential for early mouse development and acts in the cytoplasm to maintain ES cell pluripotency. Genes Dev, 2010. 24(24): p. 2772–7.

82. Wongtawan, T., et al., Histone H4K20me3 and HP1α are late heterochromatin markers in development, but present in undifferentiated embryonic stem cells. Journal of Cell Science, 2011. 124(11): p. 1878–1890.

83. Sidoli, S., et al., Metabolic labeling in middle-down proteomics allows for investigation of the dynamics of the histone code. Epigenetics Chromatin, 2017. 10(1): p. 34.

84. Sidoli, S., et al., A mass spectrometry-based assay using metabolic labeling to rapidly monitor chromatin accessibility of modified histone proteins. Scientific Reports, 2019. 9(1).

85. Wang, Z., et al., Combinatorial patterns of histone acetylations and methylations in the human genome. Nature Genetics, 2008. 40(7): p. 897–903.

86. Luo, M., et al., H3K18ac Primes Mesendodermal Differentiation upon Nodal Signaling. Stem Cell Reports, 2019. 13(4): p. 642–656.

87. Mostaqul Huq, M.D., et al., Suppression of receptor interacting protein 140 repressive activity by protein arginine methylation. The EMBO Journal, 2006. 25(21): p. 5094–5104.

88. Iwasaki, H., et al., Disruption of Protein Arginine N -Methyltransferase 2 Regulates Leptin Signaling and Produces Leanness In Vivo Through Loss of STAT3 Methylation. Circulation Research, 2010. 107(8): p. 992–1001.

89. Shen, L., et al., Loss-of-function mutation in PRMT9 causes abnormal synapse development by dysregulation of RNA alternative splicing. Nature Communications, 2024. 15(1).

90. Perez-Riverol, Y., et al., The PRIDE database resources in 2022: a hub for mass spectrometry-based proteomics evidences. Nucleic Acids Res, 2022. 50(D1): p. D543–D552.

