## Supplementary figures for "Loss of CARM1 alters the developmental programming of Glioma stem-like cells and creates a druggable NGFR/NTRK dependency"

A.

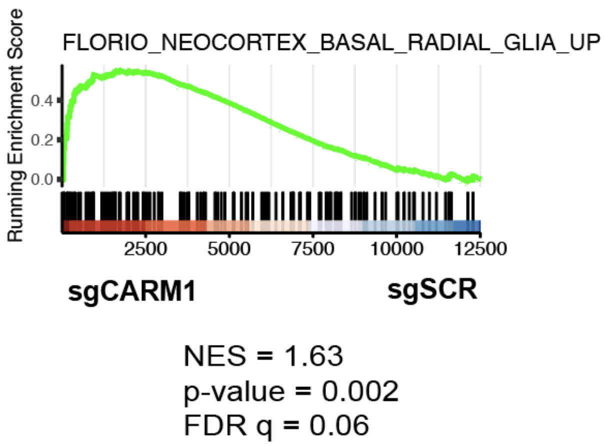


B.

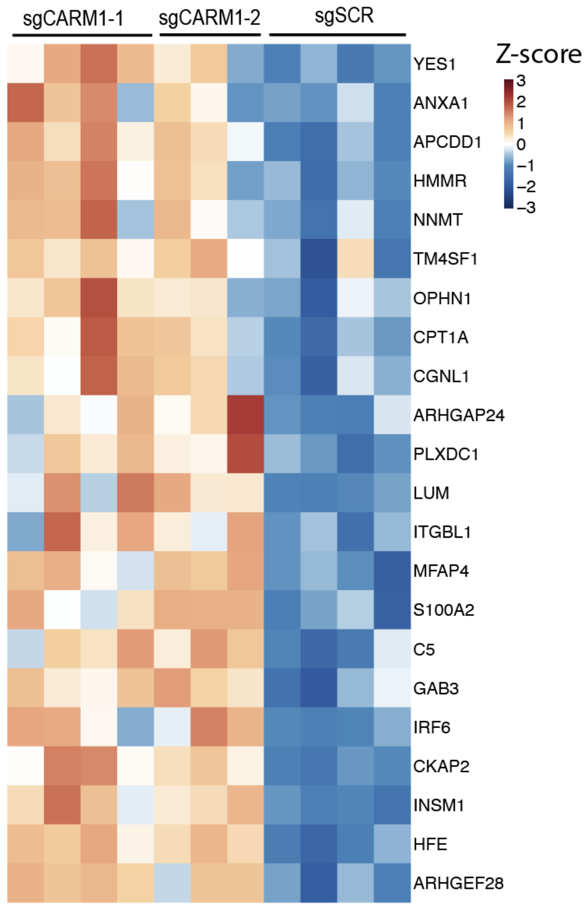


**Supplementary Figure 1.** A) GSEA of RNA-seq and B) heatmap of differentially expressed genes found enriched in radial glial cells from panel A.

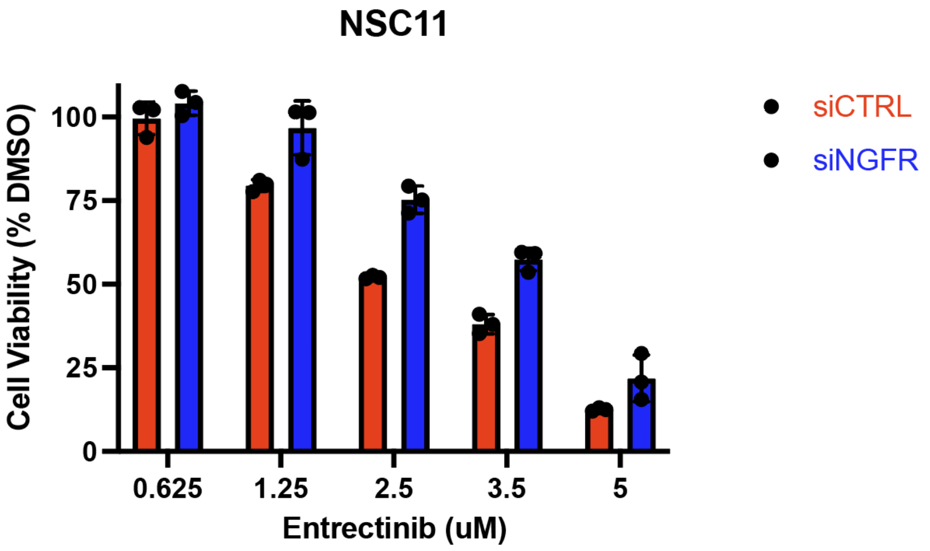


**Supplementary Figure 2.** Bar graphs after 4-day treatment of Entrectinib on siCTRL or siNGFR NSC11 GSCs, n = 3, data shows mean +/- SD.

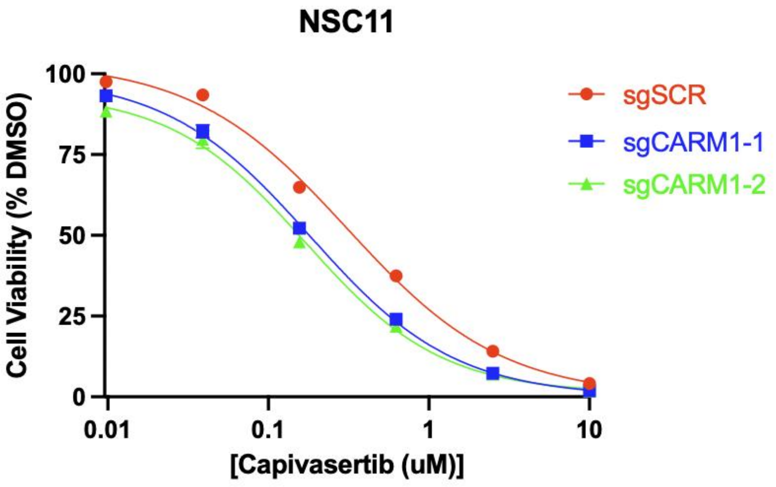


**Supplementary Figure 3.** Dose response curve of NSC11 sgSCR or sgCARM1 cells treated with the pan-AKT inhibitor, Capivasertib, n = 3. Data shows mean +/- SEM. **sgSCR ic50 = 312nM sgCARM1-1 ic50 = 182nM sgCARM1-2 ic50 = 171nM**
